# Basal p53 activity is differentially impacted by individual mutations, defining the immediate response to genotoxicity

**DOI:** 10.1101/2024.09.05.611436

**Authors:** Maria José de la Concha, Patricia Eror Barnes, Kioko Mwikali, Bee Ling Ng, Baisakhi Mondal, Mukund Khatavkar, Hannes Ponstingl, Alena Pance

## Abstract

Genotoxicity induces p53 to coordinate a cellular response, however basal p53 activity preserves genome integrity by maintaining low expression of its target genes that scan the DNA and quickly repair damage. Acquired p53 mutations disrupt this regulation and the cellular capacity to respond effectively to environmental insults. Three breast cancer cell lines with different p53 status: wild type (MCF-7) and mutated in the DNA binding domain (R280K, MDA-MB231) and (L194F, T47D) are used to identify an unbiased high-confidence subset of direct p53-target-genes. Expression of these genes is differentially impacted by the mutations in resting conditions, with a strongly reduced overall p53 activity in MDA-MB231 and mildly in T47D. Short exposure to 4NQO is used to induce DNA adducts without p53 activation as confirmed by limited changes in p53-target-gene expression. MDA-MB231 show dysfunctional DNA repair and G1 checkpoint while T47D maintain some DNA repair capacity. Both mutant lines have increased survival to genotoxicity, particularly T47D, compared to MCF-7, consistent with a different effect on p53 function and cell homeostasis. Clinical data for mutations in these residues suggests that in the balance of tumour progression impaired DNA repair is more important than proliferation and these pathways are differentially affected by specific mutations.

## INTRODUCTION

The cellular response to environmental cues is effected by appropriate gene expression controlled by activated transcription factors. The major orchestrator of the response to damaging insults is p53 that regulates critical genes involved in DNA repair, cell cycle and cell fate (Hernández Borrero and El-Deiry, 2021). As a transcription factor, it has a central DNA-Binding-Domain (DBD) with specificity for the Response Element (RE) sequence CATGCCCAGACATG, flanked by transactivation (TAD1/TAD2), oligomerisation (OD), proline-rich (PRD) and C-terminal (CTD) domains (Andreas C. Joerger and Fersht, 2010; Brady and Attardi, 2010) In steady-state p53 is maintained at very low levels due to its short half-life and thermodynamic instability, rapidly unfolding at body temperature which results in inefficient DNA binding (Lakin and Jackson, 1999). Additionally, p53 is constantly marked for degradation by the ubiquitin ligase MDM2 through interaction with TAD1(Brummer and Zeiser, 2024). DNA damage leads to phosphorylation of MDM2 that interferes with TAD1 binding thereby increasing p53 levels. Post-translational modifications stabilise and activate p53 which tetramerises on the p53-REs in target gene (TG) promoters (Abuetabh *et al*., 2022). The TGs halt cell cycle progression, coordinate DNA damage repair (DDR) and if it fails, trigger apoptosis. However, basal p53 activity maintains low expression of critical TGs for background scanning of naturally-occurring common damage to maintain genomic integrity and protect cells from intrinsic and environmental stress (Loewer *et al*., 2010; Seim, Ma and Gladyshev, 2016; Pappas *et al*., 2017; Kotler *et al*., 2018). Many of the effector genes are induced directly by p53 binding to their promoters but others are controlled by some of the prominent p53-TGs, such as *CDKN1A* (p21)(Engeland, 2022).

Downregulation of p53 expression by knock-out or dysregulation by missense mutations, particularly in the DBD, leads to decreased levels of p53-TGs and has been correlated with tumorigenesis (Lane, 1992; Levine, 1997). Unsurprisingly, the gene that encodes p53 (*TP53)* is the most frequently mutated in human cancer (Baker *et al*., 1989; Nigro *et al*., 1989; Kennedy and Lowe, 2022), and breast cancer is no exception (Wasielewski *et al*., 2006), constituting the highest cause of female mortality (Bray *et al*., 2024). The mainstream treatments are chemo and radiotherapy (Markowska *et al*., 2024), that generally aim to induce cell death by damaging the DNA. This response depends largely on p53 and mutations that impair its activity have profound impact on the outcome of therapy. While mutations have been detected all along the *TP53* gene (Bouaoun *et al*., 2016; Tate *et al*., 2019) most occur in the DBD (Kotler *et al*., 2018) impacting the stability and capacity to bind the DNA REs (Kato *et al*., 2003). It has been shown that some regions of the p53 DBD are fairly permissive to aminoacid variation while others are highly intolerant to change. Consequently, distinct mutations affect p53 function differently either by decreasing DNA binding efficiency or modulating interactions with other proteins (Kotler *et al*., 2018), with an impact on pathway activation (Daily *et al*., 2011). Understanding the regulatory landscape of p53 and the impact different mutations have on its activity could have great implications for therapy design. Deciphering p53 basal activity will contribute to understanding the mechanisms of p53 and the role of spontaneous somatic mutations in the protein in tumorigenesis.

This work uses three breast cancer cell lines with different p53 status: wild type MCF-7 and two with non-hot spot mutations in the DBD: MDA-MB231 (R280K) and T47D (L194F), to identify direct p53-TGs and understand basal p53 function. Transcriptomics and ChIP-seq data were combined with existing sources to generate a list of high-confidence direct p53-TGs. DNA lesions were induced with short exposure to 4NQO to study the immediate response that relies on basal p53 activity and how it is affected by different mutations in the DBD. Combining cellular assays and transcriptomics we show that R280K causes a dramatic decrease of overall p53 activity characterised by failure to repair DNA adducts and to arrest in G1. L194F shows a milder effect, retaining some DNA repair capacity. Both mutants have increased survival compared to MCF-7, demonstrating that p53 basal activity that maintains low levels of crucial TGs is critical for the rapid response to genotoxicity. The implications for disease outcomes are assessed using cancer data bases to analyse tumours with mutations in these residues. The clinical data shows marked decrease in survival in both cases but distinct mutations seem to affect outcomes differently, which could have implications for prognosis and therapeutic choices. Understanding the mechanisms of p53 and the impact of different mutations can contribute to the management of the disease.

## RESULTS

### A comprehensive list of direct p53 target genes shows differences in expression in resting cells with different p53 status

Three breast cancer cell lines with distinct characteristics (Table I) are used to explore the basal functionality of p53-regulated cellular processes. The MCF-7 line expresses a wild type (WT) p53 while MDA-MB231 and T47D present mutations in the *TP53* DBD. The mutations are located in different regions of the DBD: MDA-MB231 R280K and T47D L194F and do not have a major impact on p53 expression (Fig.1A).

**Table I.** Characteristics of the breast cancer cell lines MCF-7, MDA-MB231 and T47D. Information obtained from ATCC, Cellosaurus(Bairoch, 2018) and Cosmic(Tate et al., 2019).

| Cell Line | Tumor Type | MOLECULAR CLASSIFICATION |  | RECEPTOR CHARACTERISTICS |  |  |  |  |  |  |
| --- | --- | --- | --- | --- | --- | --- | --- | --- | --- | --- |
|  |  | General Subtype | Subgroup | ER (*ESR1*) | PR (*PGR*) | HER2 (*ERBB2*) | EGFR | p53 (*TP53*) | BRCA1 | BRCA2 |
| MCF7 | AC | ER+ | Luminal A | + | + | ++ | - | WT | WT | WT |
| MDA-MB-231 | AC | TNBC | Basal B | - | - | + | + | R280K | WT | WT |
| T47D | DC | ER+ | Luminal A | + | ++ | ++ | - | L194F | WT | WT |

To understand the regulatory landscape of p53, it is essential to identify its directly regulated subset of genes. For this, genome-wide p53 DNA binding data was obtained from the ChIP-Atlas database (Zou, Ohta and Oki, 2024), extracting genes with peaks detected 1Kb around their transcription start sites (TSS) in breast lines and tissues. Peaks with a minimum binding score of 100 yielded a list of 9382 potential p53-TGs. A comparison with a list of 3661 genes identified by an exhaustive study from 2017 (Fischer, 2017) and a curated list of 2740 potential targets of varying confidence from the DoRothEA regulon site (Garcia-Alonso *et al*., 2019) revealed a substantial but not complete overlap of the potential direct p53-TGs. Thus, 1440 genes from DoRothEA show p53 binding in the ChIP-Atlas, which also contained 2228 genes from the 2017 study (Fischer, 2017). Based on the premises that p53 basal activity maintains low-level expression of its TGs (Loewer *et al*., 2010; Pappas *et al*., 2017) and mutations in p53 DBD interfere with promoter binding, transcriptomic profiles of these cell lines were used to perform an unbiased analysis, overlapping differentially expressed genes with the lists of TGs identified in the different sources (Fig. 1B). Differentially expressed genes present in at least two of the databases used were compiled in a list of 2052 high-confidence direct p53-TGs and their expression pattern in the cell lines is shown in Figure 1C.

**Figure 1:**
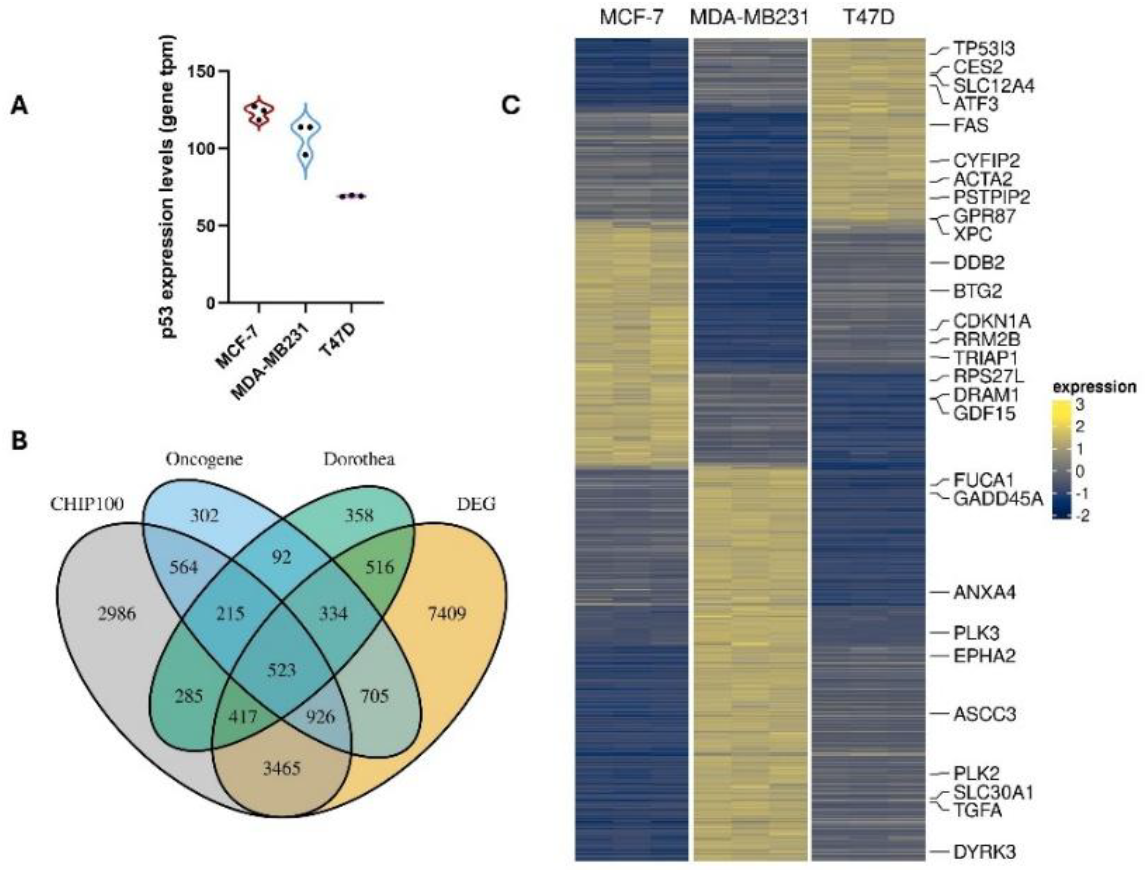
direct p53 target gene subset. **A:** expression levels of p53 in MCF-7 (WT), MDA-MB231 (R280K) and T47D (L194F) detected by RNAseq (gene tpm); **B:** Venn diagram of putative direct p53 target genes identified in different data sets, overlapped with differentially expressed genes in the three cell lines. **D:** Heatmap of high confidence direct p53 target genes differentially expressed in the three cell lines, found in at least two p53 target data bases.

The transcriptomic data of the direct p53-TGs was used to estimate overall p53 activity (Fig.2A), which revealed that the MDA-MB231 R280K mutation causes a dramatic decrease while the T47D L194F mutation has a milder effect, with an activity closer to MCF-7. The cellular pathways regulated by the p53-TGs was explored using the Reactome pathway database (Milacic *et al*., 2024) and a gene set enrichment analysis (GSEA) was performed to compare pathway expression between the cell lines (Fig. 2B, Supplementary data). Disruption of p53 transcriptional activity is evident, and interestingly, while DNA repair and cell cycle checkpoint related genes are more noticeably downregulated in MDA-MB231 than T47D, mitotic processes seem upregulated in MDA-MB231.

**Figure 2.**
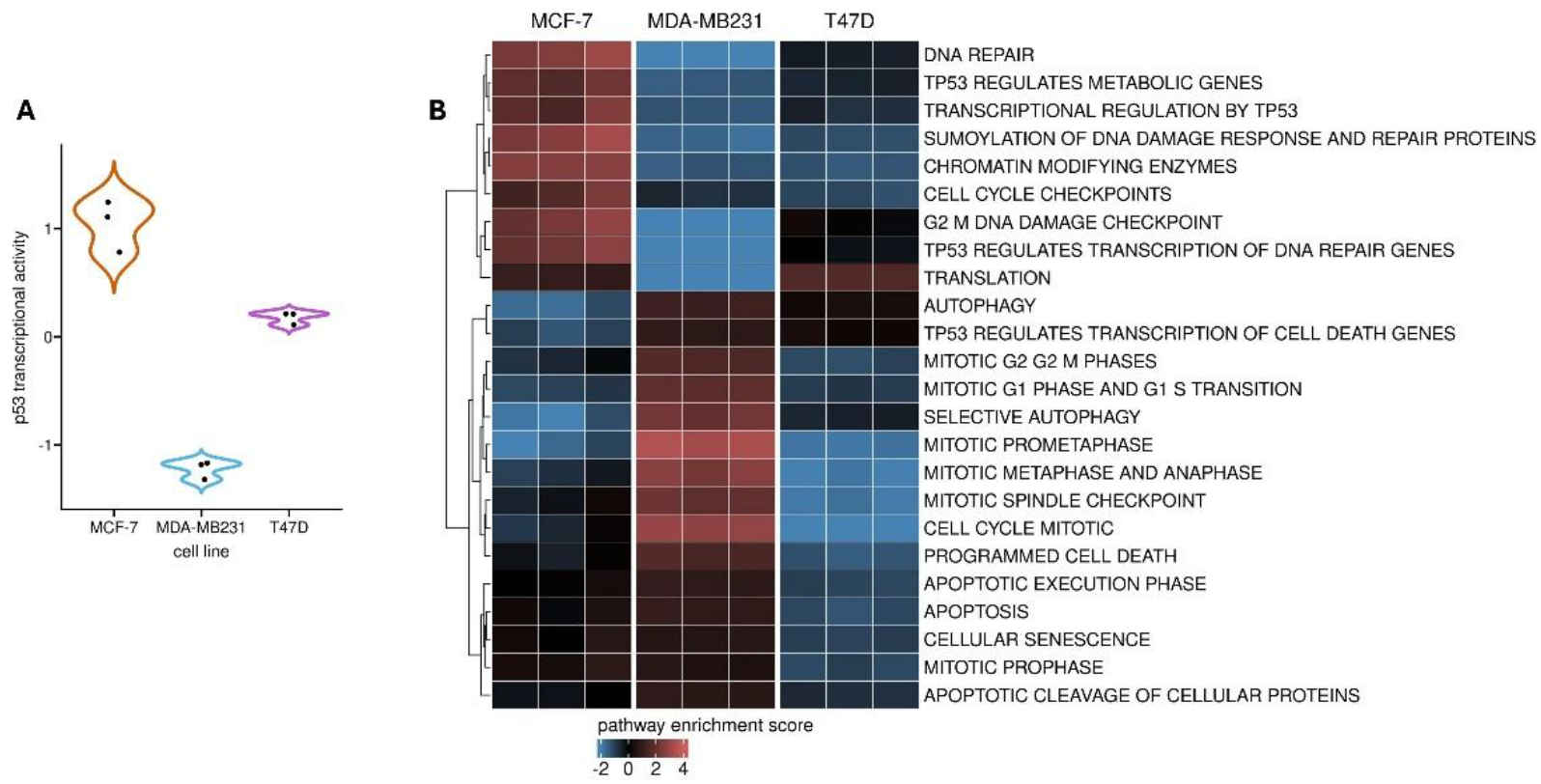
p53 differential activity in breast cancer cell lines with different p53 status. **A:** estimation of p53 activity in resting cells, using a univariate linear model and the progeny p53 markers/targets. **B:** heatmap of GSEA for expression of p53 relevant pathways according to the Reactome data base using RNAseq data from resting cells.

### The immediate response to DNA bulky lesions relies on basal expression of p53-TGs

The response to naturally occurring insults was mimicked using 4-nitroquinoline 1-oxide (4NQO) to cause DNA lesions. 4NQO is metabolised intracellularly into 4-hydroxyamino-quinoline 1-oxide (4HAQO) that forms bulky adducts on the DNA (Khiavi *et al*., 2019). These adducts form within minutes of exposure with a maximum level occurring after approximately 1 hour and are swiftly repaired mainly by the Nucleotide Excision Repair (NER) mechanism (Adar *et al*., 2016) (Fig. 3A). Excision of the damaged nucleotides during repair can lead to Single Strand Breaks (SSB) in a concentration-dependent manner. This type of damage is repaired by the Single Strand Break repair mechanism (Fig. 3A), that in normal cells can repair most SSBs within 2 hours (Lankinen, Vilpo and Vilpo, 1996). The early response to DNA lesions was studied exposing the cells to a low (1µM) and high (10µM) concentration of 4NQO for 2 hours.

**Figure 3.**
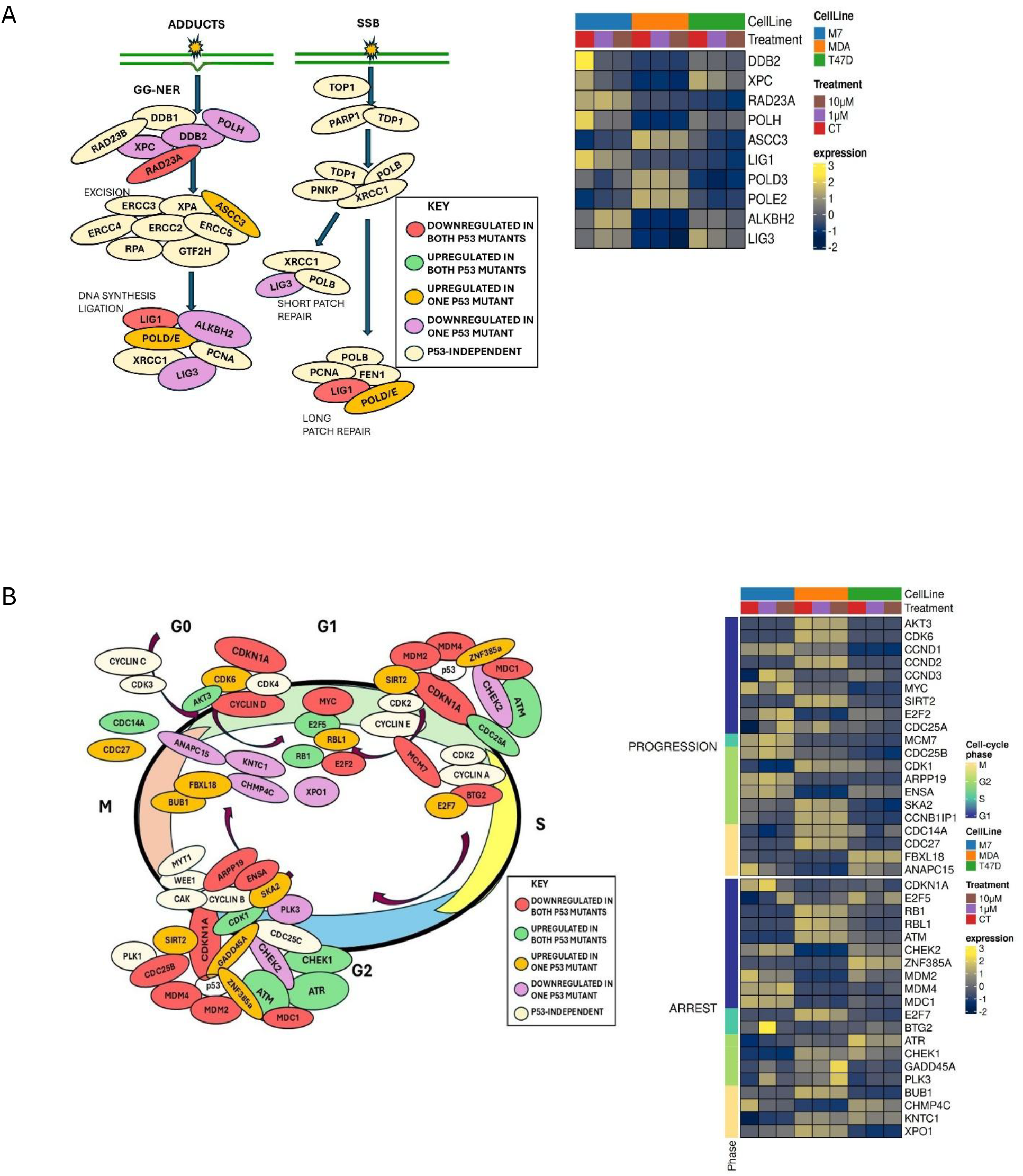
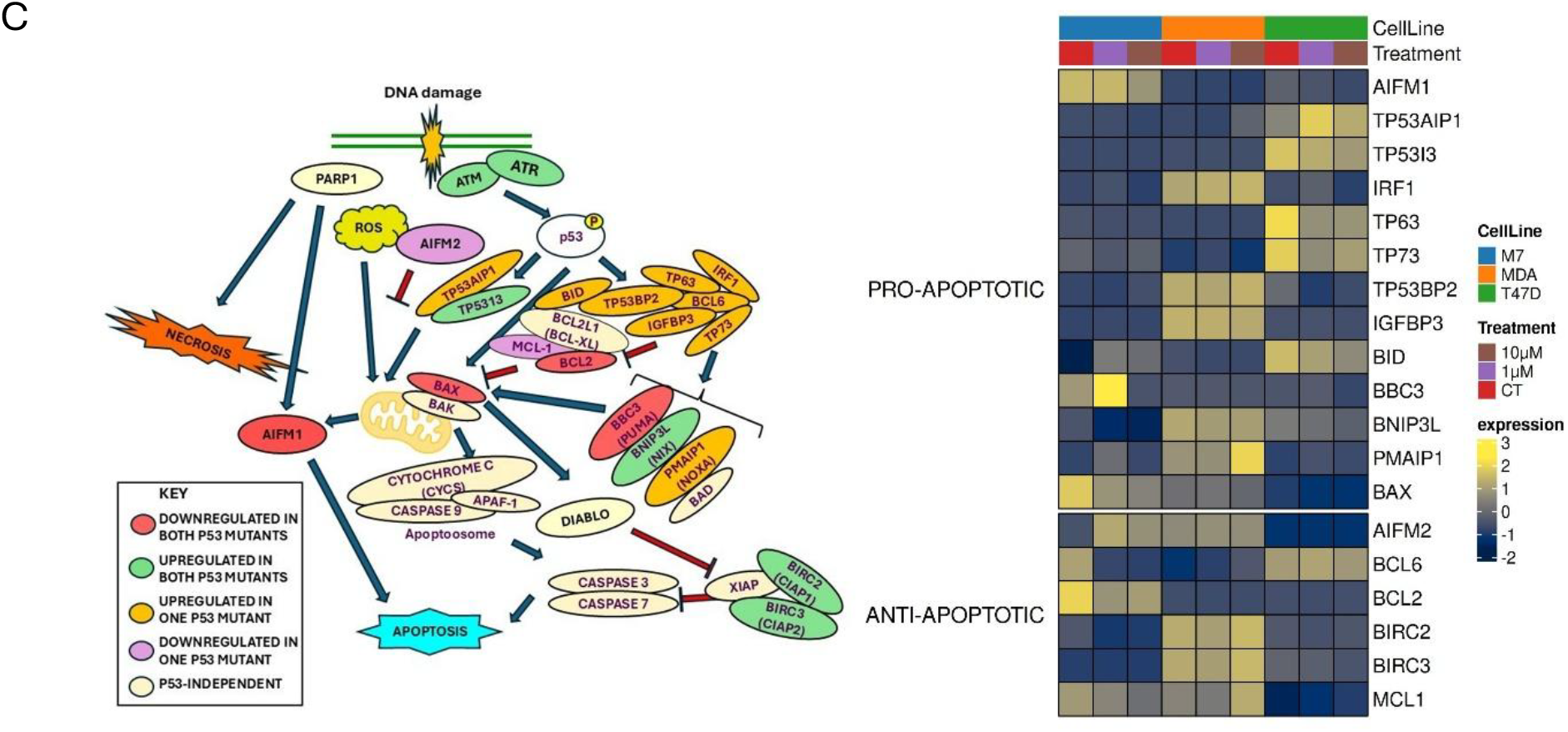
cellular pathways involved in the DNA damage response and expression of participating p53-TGs. **A) left:** cartoon of the mechanisms that repair DNA adducts and SSBs; **right:** heatmap of the expression of p53-TGs in this pathway. **B) left:** cartoon of the cell cycle regulation checkpoints; **right:** heatmap of the p53-TGs involved **C) left:** cartoon of apoptotic cell death; **right:** heatmap of the p53-TGs of this pathway. The cartoons present the genes directly involved in the regulation of each pathway and these are colour-coded to represent those genes that are up-regulated in one (orange) or both (green) and down-regulated in one (purple) or both (red) mutant cell lines. The p53-independent genes involved are cream. The heatmaps are constructed with RNAseq expression (gene tpm) and z-scored scaling.

Treatment with 4NQO caused only subtle changes in gene expression of the three cell lines. The p53-dependent genes involved in NER are mostly downregulated in MDA-MB231 (purple) or both (red) mutant cell lines (Fig. 3A heatmap). Despite a slight increase in expression in some of these genes (*POLD/E, ASCC3*) and steady or increased expression of the p53-independent genes involved in this pathway (Fig. S3), the critical adduct recognition genes, *XPC* and *DDB2* (Gunz, Hess and Naegeli, 1996) are downregulated in MDA-MB231 (Fig. 3A). This suggests that NER is impaired in MDA-MB231 but active at least to some extent in T47D. Interestingly, T47D expression of *PolH*, capable of bypassing DNA adducts (Klarer *et al*., 2012), is similar to WT p53 MCF-7, while it is greatly downregulated in MDA-MB231 (Fig. 3A).

The p53-TGs involved in the regulation of the cell cycle (Fig. 3B, expression of p53-independent genes involved in cell cycle control are presented in Fig. S4) are mostly downregulated in both mutant lines (red). However, some of them are upregulated in MDA-MB231 in these conditions when p53 has not yet been activated (Fig. 3B heatmap). Notably, the regulatory axis *AKT3-CDK6-CCND1*(Cyclin D1), that can promote cell cycle progression, particularly through G1 are clearly expressed in this cell line (Fig. 3B).

Examination of the intrinsic programmed cell death pathway (Fig. 3C) shows that many of the pro-apoptotic p53-TGs are upregulated in the mutant cell lines (Fig. 3C, heatmap). Particularly, the critical gene for the initiation of apoptosis *BAX* and *AIFM1* that can induce apoptosis in a p53-dependent and independent fashion are still expressed (Fig. 3C). At the same time, *BCL6* levels are high in T47D and the p53-independent anti-apoptotic *BCL2L1* (BCL-xL) is much higher in both mutant cell lines (Fig. S5).

### Basal p53-coordinated response to short-term DNA adducts is differentially affected by specific mutations in the DBD

The immediate response to 2h 4NQO exposure is a small decrease of cell density at both low (1µM) and high (10µM) concentrations, that only becomes significant in MDA-MB231 cells (Fig. 4A). Because the replication time of these cell lines is *ca*. 24 hours, the effect of 4NQO on cell homeostasis was assessed by returning cells to fresh medium for 24h after treatment. The recovery period shows a significant fall in cell density in MCF-7 at both concentrations while MDA-MB231 and T47D maintain similar density levels to those immediately after treatment.

**Figure 4.**
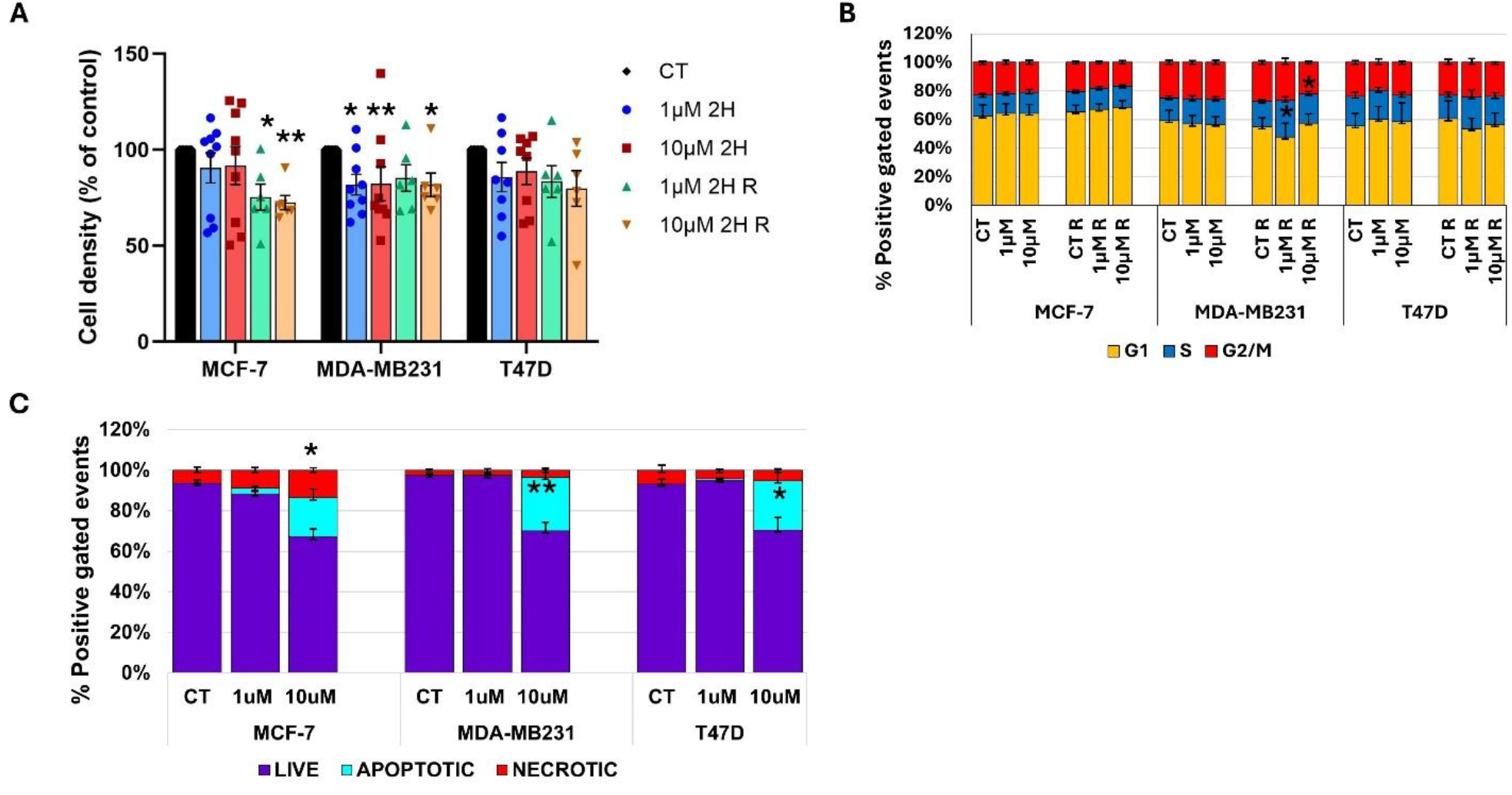
Immediate response of the major cellular pathways to genotoxic insult. Cells were treated with 4NQO 1µM and 10µM for 2h. **A)** Cell density measured by MTTA after treatment and cells recovered for 24h after removal of 4NQO. Mean and SEM of at least 8 biological replicas. **B)** Cell cycle phases measured by flow cytometry of cells stained with propidium iodide of treated and rescued cells. Mean and SEM of at least 4 biological replicas. **C)** Cell death caused by 4NQO measured by flow cytometry labelling necrotic cells with propidium iodide and apoptotic cells with Hoechst33342. Mean and SEM of 4 biological replicas. *P<.05, **P<.01

The regulation of the cell cycle showed little change with only subtle tendencies of concentration-dependent G1 retention in MCF-7 and T47D indicating a delay of S entry, consistent with DNA replication stalling to allow adduct repair (Fig. 4B). MDA-MB231 appear to be ‘frozen’ with almost no change. The recovery follows the same trend, though significant changes occur in MDA-MB231, namely an increase of the S population after 1µM and a decrease in G2/M at 10µM, reflecting impaired DNA repair mechanisms that slow down DNA replication and dysfunctional cell cycle control. T47D also show an expansion of the S population though it is not significant.

Cell death following short exposure to 4NQO revealed a slight non-significant increase in MCF-7 while negligible in the mutant lines after 1µM 4NQO. This is consistent with a delay in the cell cycle to allow DNA repair, yet the significant decrease in cell density in MDA-MB231 indicates a complete cell cycle arrest correlated with non-functional NER (Fig. 4C). At 10µM 4NQO the intensity of the damage seems to overwhelm the repair machinery triggering a significant rise in necrosis and apoptosis in MCF-7 cells. The mutant cell lines on the other hand, show a significant increase in apoptosis, while necrosis remains negligible. Interestingly, the total proportion of cell death is very similar in all cell lines and yet the mutant lines show a remarkable recovery of cell density after 24 hours, particularly MDA-MB231 which showed a significant drop after treatment.

### Survival outcomes in breast cancer patients with mutations in the R280 and L194 residues in the p53 DNA Binding Domain

Clinical profiles from The Cancer Genome Atlas (TCGA) (Koboldt *et al*., 2012), the Molecular Taxonomy of Breast Cancer International Consortium (METABRIC) (Curtis *et al*., 2012) and the Memorial Sloan Kettering Cancer Center (MSK) (Zehir *et al*., 2017) were explored via the cBioPortal platform (Cerami *et al*., 2012) to gather clinical data from breast cancer patients with wild type p53 and mutations in the residues analysed here. The numbers are low and the specific L194F substitution was not found. Nevertheless, cases with mutations in L194, R280 and the specific R280K were gathered and survival outcomes were compared to wild type p53 (Table II).

**Table II:** survival outcomes of patients with breast tumours carrying mutations in the p53 residues R280 and L194 compared to WT p53. Information obtained from TCGA, METABRIX and MSK.

|  | WT P53 | % | P53 R280 (S,T,G,I) | % | P53 R280K | % | P53 L194 (ALL) | % |
| --- | --- | --- | --- | --- | --- | --- | --- | --- |
| **TOTAL SAMPLES** | 918 |  | 39 |  | 22 |  | 48 |  |
| **DECEASED** | 41 | 4.4 | 20 | 51.3 | 11 | 50 | 22 | 45.8 |
| **LIVING** | 878 | 95.6 | 19 | 48.7 | 11 | 50 | 26 | 54.2 |
| **SURVIVAL TIME** |  |  |  |  |  |  |  |  |
| **UP TO 1 YEAR** | 134 | 14.6 | 8 | 20.5 | 1 | 4.5 | 6 | 12.5 |
| **1 TO 5 YEARS** | 584 | 63.6 | 16 | 41 | 14 | 63.6 | 30 | 62.5 |
| **OVER 5 YEARS** | 214 | 23.3 | 15 | 38.5 | 7 | 31.8 | 12 | 25 |
| **TMB MEDIAN** | 1.3 |  | 2.5 |  | 5.5 |  | 3.6 |  |

TMB: tumour mutation burden

As expected, mutation of both residues dramatically impact survival with L194 showing slightly better outcome than R280, in agreement with the critical role of R280 for p53 function. The higher mutation burden reflects the ineffective DNA repair caused by p53 dysfunction and particularly by the R280K substitution. Surprisingly, survival time is similar between the wild type and L194 groups while the R280 mutation seems more aggressive, causing higher death rates within a year but paradoxically, long-term survival (>5 years) is slightly better.

Separation of the cases by substitution shows not all are present and no clear prevalence of 1 vs 2 nucleotide changes is observed (Table III). All the L194 all the substitutions found are 1 nucleotide changes and the 2 nucleotide change F is absent, while all R280 substitutions are 2 nucleotide changes except G which is not the most common.

**Table III:** survival outcomes of patients with breast tumours with specific substitutions in the mutated residues of p53. Information obtained from TCGA, METABRIX and MSK

| mutated amino acid | R280 |  |  |  |  |  |  | L194 |  |  |  |  |
| --- | --- | --- | --- | --- | --- | --- | --- | --- | --- | --- | --- | --- |
| specific substitution | K | % | S | T | % | G | I | R | % | H | P | % |
| total samples | 25 |  | 4 | 26 |  | 5 | 8 | 39 |  | 1 | 14 |  |
| samples with known outcome | 22 |  | 3 | 22 |  | 6 | 8 | 35 |  | 1 | 12 |  |
| deceased | 11 | 50 | 0 | 13 | 59 | 3 | 4 | 17 | 49 |  | 5 | 41.7 |
| alive/ tumour free | 11 | 50 | 3 | 9 | 41 | 3 | 4 | 18 | 51 | 1 | 7 | 58.3 |
| SURVIVAL |  |  |  |  |  |  |  |  |  |  |  |  |
| UP TO 1 YEAR | 1 | 4.5 |  | 7 | 32 | 1 | 0 | 5 | 14 |  | 1 | 8.3 |
| 1 TO 5 YEARS | 14 | 64 |  | 5 | 23 | 2 | 6 | 19 | 54 | 1 | 10 | 83.3 |
| OVER 5 YEARS | 7 | 32 |  | 10 | 46 | 3 | 2 | 11 | 31 |  | 1 | 8.3 |

The R280 mutation has lower survival than L194 again consistent with stronger effect on p53 activity. The survival time is generally better for R280, but the trend of aggressivity and long-term survival is more prominent in the T substitution. This shows that not only the residue affected is important but also the specific substitution has a different impact on outcome.

## DISCUSSION

The function of p53 largely depends on promoter binding through REs, therefore it is not surprising that most mutations affecting its function are found in the DBD spanning amino acids 94-292 (Walerych *et al*., 2012; Bouaoun *et al*., 2016). To understand p53 function and the impact of these mutations, it is critical to identify the subset of direct p53-TGs. This is not trivial due to the technical challenges of the mainstream approaches used, such as antibody specificity, illustrated by the differences in the p53-TGs reported in previous studies (Fischer, 2017; Garcia-Alonso *et al*., 2019). Additionally, the broad regulatory function of p53 translates in a high number of responsive genes, many of which are indirectly modulated through intermediary proteins. Basal p53 activity maintains expression of its TGs (Loewer *et al*., 2010; Pappas *et al*., 2017) and we show that this is reflected in the expression levels of some of the genes involved in p53-dependent pathways. Based on this observation combining transcriptomics of p53-defective cell lines with existing ChIP data bases and published resources is an unbiased approach that generated a high-confidence list of p53-TGs.

The expression profiles of the p53-TGs reveal a different impact of the two DBD mutations examined on basal p53 activity, in agreement with previous reports of differential effects of p53 mutations (Walerych *et al*., 2012; Kotler *et al*., 2018). The dramatic decrease in the overall p53 activity caused by the MDA-MB231 R280K mutation is consistent with a fundamental role in DNA binding (Gomes *et al*., 2018). This arginine forms hydrogen bonds with the highly conserved guanine (CAT**G**TCC) in the RE and with a structural molecule of water and surrounding residues Cys277 and Asp281. It also forms a stabilising salt bridge network with Asp273 to establish sequence-specific contacts in the major groove (Andreas C Joerger and Fersht, 2010). Indeed, a p53 variant lacking residues 257-322 is incompetent for DNA binding and unable to induce p21(García-Alai *et al*., 2008). On the other hand, the mutated residue in T47D (L194), forms part of the L2 loop of the DBD that, while involved in DNA binding (Young *et al*., 2008) is one of the regions more tolerant to alterations (Kato *et al*., 2003; Kotler *et al*., 2018). This is consistent with the milder impact of this mutation on p53 activity. Interestingly, many p53-TGs are upregulated in the mutant lines compared to MCF-7, which could indicate a repressive activity of p53, though the general thinking is that p53 is exclusively an activator (Fischer, 2017). Nevertheless, some studies have shown repressive activity (Allen *et al*., 2014) and others have reported a gain-of-function effect by certain mutations (Bargonetti and Prives, 2019; Butera and Amelio, 2025), yet this effect has not been documented for either of the mutations studied here. Alternatively, it is possible that these genes are co-regulated by p53 with other TFs and therefore respond differently to p53 inactivation.

The limited effect of genotoxic damage caused by short exposure to 4NQO on p53-TGs expression is consistent with p53 activation not being triggered. This is expected because the bulky adducts caused by 4NQO are repaired by basal NER (Adimoolam and Ford, 2003), as they are not recognised by the major p53-activating proteins ATR and ATM (Sugasawa, 2016). NER depends on constant scanning of the genome for DNA lesions by basal expression of p53-TGs such as *XPC, DDB2* (Hwang *et al*., 1999; Wang *et al*., 2003) and *Rad23*. The damage is verified by XPA and RPA, helicases unwind the DNA, the adducts are excised by the ERCC complex (Choi *et al*., 2014) and the gap is rapidly filled by DNA polymerases. The transcriptome data shows lower levels of the p53-TGs involved in sensing DNA modification indicating that NER is dysfunctional in MDA-MB231 while only mildly affected by the T47D mutation. This is confirmed by the very slight (non-significant) decrease in cell density in MCF7 and T47D because repairing of the DNA adducts allows quick reset of the cell cycle and consistent with little change in the cell cycle phases. Additionally, both express *POLH* which can bypass the adducts and allow DNA replication to proceed (Lange, Takata and Wood, 2011; Dhoonmoon *et al*., 2025). On the other hand, the significant decrease in MDA-MB231 cell density, accompanied by a static cell cycle vouches for stalled proliferation. The lack of cell death in all cell lines after 1µM 4NQO is consistent with delayed DNA replication. Any SSBs formed at this time are quickly recognised by Parp1 that recruits enzymes to process the broken ends (*PNKP, ATPX*), fill the gap (*POLB*) and ligate the strand (*Lig1* and *3*) in a largely p53-independent fashion. However, the DNA lesions caused by 4NQO are concentration-dependent and at 10µM the level of damage overwhelms the repair machinery, producing higher levels of SSB (Fujiwara, 1989), that lead to replication fork stalling and collapse leading to Parp1 activation and possibly ROS formation. Parp1, which is upregulated in both mutant cell lines, causes mitochondrial outer membrane (MOM) permeabilization (Yu *et al*., 2006) with the consequent release of AIF (*AIFM1*) into the cytoplasm. AIF translocates into the nucleus where it initiates apoptosis through chromatin condensation and DNA fragmentation in a p53 and caspase-independent fashion (Cheung *et al*., 2005). The apoptotic process can lead to secondary necrosis (Silva, 2010), consequence of membrane compromise resulting from cellular stress. The lower basal levels of *AIFM1* in the mutant lines lead to slower apoptosis and as a consequence, low levels of secondary necrosis compared to MCF-7. It is intriguing that despite the increase in cell death, there is no further decrease in cell density at this concentration, which reflects very early apoptosis when cells are still metabolically active and attached.

Assessment of survival capacity in the recovery period shows a marked decrease in cell density in MCF-7 without the onset of cell death at 1µM 4NQO, indicating a slow reset of the cell cycle despite effective DDR. The evident expansion of the S population in the mutant lines shows ineffective G1 checkpoint and a struggle to replicate due to faulty DDR, particularly in MDA-MB231. Accordingly, *CDKN1A* (p21) is more prominently downregulated in MDA-MB231 and T47D compared to MCF-7. The increased S population in MDA-MB231 could be correlated with overexpression of *AKT3* and *CDK6* which promote G1-S transition (Tsai *et al*., 2022). Indeed, Akt3 has been proposed as a therapeutic target in triple-negative breast cancer (Chin *et al*., 2014). After recovery from 10µM 4NQO, cell density also decreases significantly reflecting the consequences of high levels of cell death caused by overwhelming DNA damage. The persistent DNA damage triggers p53 activation throughout the recovery period via the ATR-H2AX-CHEK1 pathway that recognises SSBs and ATM-DNA-PK-CHEK2 that recognises DSBs and induces HR and NHEJ DNA repair mechanisms. Intriguingly, in contrast to MCF-7, cell density after recovery in the mutant lines is similar to just after treatment, showing high survival capacity of these cells. This could be related to sustained expression of anti-apoptotic BCL6, which has been associated with oncogenicity and resistance to genotoxic stress (Wu *et al*., 2014; Liu *et al*., 2022).

Clinical data aligns with the critical nature of R280 for p53 function as we show in the model cell line MDA-MB231. While the R280K substitution causes higher mutation burden, possibly correlated with its dysfunctional DDR, it is R280T that seems more aggressive. The higher long-term survival of patients with mutations in R280 despite the overall worse outcomes suggests that the critical factor is DDR. Thus, survival timeline of patients with mutations in L194 is similar to those carrying a WT p53, yet the T47D line shows higher proliferation and survival but maintains DNA repair capacity. Perhaps on the longer term, mutation accumulation results in reduced fitness. Mutations in p53 worsen disease outcomes but the impact of specific residues and substitutions is slightly different. In the balance between cell cycle and DNA repair dysregulation that will determine tumour progression, DNA repair seems more important for a better prognosis. It should be taken into account that the numbers of patients with these mutations are small and there are additional factors involved such as treatment and genomic background of the patients that might include mutations in other important genes with impact on the disease.

The distinct effect of particular mutations in p53 on its activity and the cellular response to genotoxicity highlights the need for an in-depth mechanistic understanding of their impact. This could represent a useful molecular tool for therapy choice, prediction of tumour behaviour as well as prognosis and therapy outcomes.

## METHODS

### Cell culture

Breast cancer cell lines MCF-7. MDA-MB231 and T47D were maintained in Dulbecco’s modified Eagle’s medium high glucose (GIBCO) supplemented with 10% foetal calf serum (Fisher Scientific), 2% glutamine (GIBCO) and 1% penicillin/streptomycin (Sigma/Merck) and cultured at 37^0^C in 5% CO_2_. DSB were induced with 4-nitroquinoline 1-oxide (4NQO) (Sigma/Merck).

### RNA extraction and sequencing

Cells were treated with 1µM and 10µM 4NQO for 6 hours and total RNA was extracted with Isolate II RNA mini kit (Bioline) following the manufacturer’s instructions. Ribominus rRNA depletion kit (ThermoFisher) was used enrichment of messenger RNA. RNA quantification and quality was assessed using Bioanalyzer Pico RNA chips and cDNA were synthesised using Superscript III reverse transcriptase (Invitrogen) following the manufacturer’s instructions. cDNA libraries were prepared with NEBNext Ultra II DNA library Prep Kit (New England Biolabs) following the kit protocol and sequenced on Illumina NovaSeq within the Wellcome Sanger pipelines.

### MTT assay

Cells were seeded in 96 well plates at 5000 cells per well and left to attach over-night. 24-hour post-seeding, 4NQO was added at 1µM or 10µM for 2 hours or 24 hours. Cells were either measured for viability with an MTT assay or washed with PBS (GIBCO), replacing fresh medium for 24 hours of recovery time and cell viability was measured. Twenty µl of 5mg/ml MTT (3-(4,5-dimethylthiazol-2-yl)-2,5-diphenyltetrazolium bromide) were added per well and incubated at 37^0^C for 3 hours to form formazan crystals. The reagent and media were removed, and the formazan crystals were then solubilised in 70% ethanol for 30 minutes. Absorbance was measured at 570nM in a plate reader. Absorbance of 4 to 6 replicates was averaged and normalised to controls.

### Flow cytometry

Cells were seeded in 6 well plates and left to attach over-night. After treatment with 4NQO, the cells were carefully lifted off the plates by trypsinisation and processed depending on the downstream protocol. Samples were measured on a BD LSR Fortessa™ cell analyser (BD Biosciences).

### Cell death assessment

live cells were stained for detection of apoptosis with Hoechst-33342 (excitation/emission: 352/454nm) and necrosis with Propidium Iodide (excitation/emission: 535/615nm). Briefly, cells were resuspended in cold PBS containing 0.5µg/ml Hoechst-33342 and 0.1µg/ml and analysed by flow cytometry. The gating strategy is shown in Figure S1: the cell population was chosen on SSC-A/FSC-A eliminating debris (A), from this population doublets were excluded using FSC-W/FSC-A (B) and the singlets population was examined using the 561nm laser 610/20 filter for PI and the 405 laser 450/50 filter for Hoechst-33342. Analysis of unstained, single stained and untreated cells was used to establish the quadrant gates (C), which allowed quantification of the apoptotic cells (high intensity HO staining) in the bottom right quadrant, and the necrotic cells (positive for PI) in the top two quadrants (D). High intensity HO staining reflects nuclear condensation characteristic of apoptosis, while cells positive for PI have compromised membranes, which correspond to necrosis.

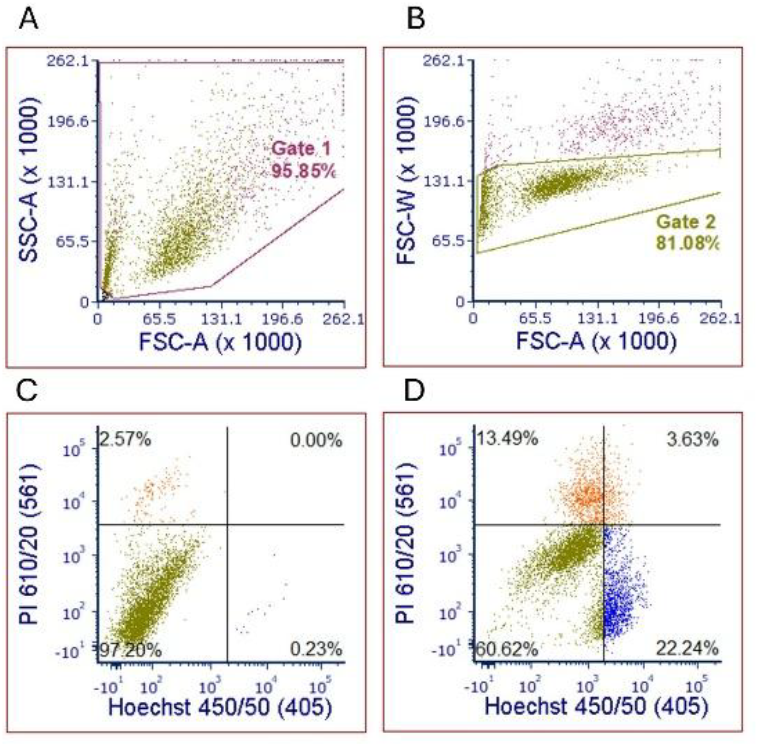

### Cell cycle analysis

cells were recovered in cold ethanol (70% in PBS) and fixed over-night at 4^0^C. Samples were then centrifuged at 290xg for 5 minutes. The ethanol was discarded, cells were washed 1x with PBS and resuspended in the staining solution: PBS with tritonX-100 (0.1%), RNAse A (SIGMA R6513) (50µg/ml) and Propidium Iodide (25µg/ml). The samples were left staining over-night at 4^0^C and analysed the following day using the 561nm laser 610/20 filter on a linear scale. The gating strategy for the cell cycle is shown in Figure S2. As above, debris is eliminated from analysis on SSC-A/FSC-A (A) and doublets excluded on PI-W/PI-A (B). The resulting population was analysed for PI fluorescence and the cell cycle phases observed and quantified in a histogram (C) according to intensity reflecting G1 (2n), S (duplicating DNA) and G2/M (4n).

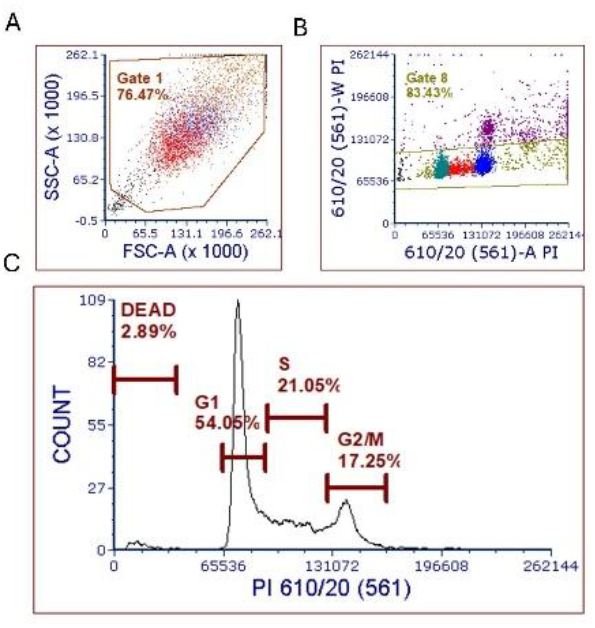

## STATISTICAL ANALYSIS

Significance of the differences in cell behaviour detected with the different techniques was assessed by an unpaired two two-tailed distribution T-test.

## SUPPLEMENTYARY FIGURES

**Fig. S3.**
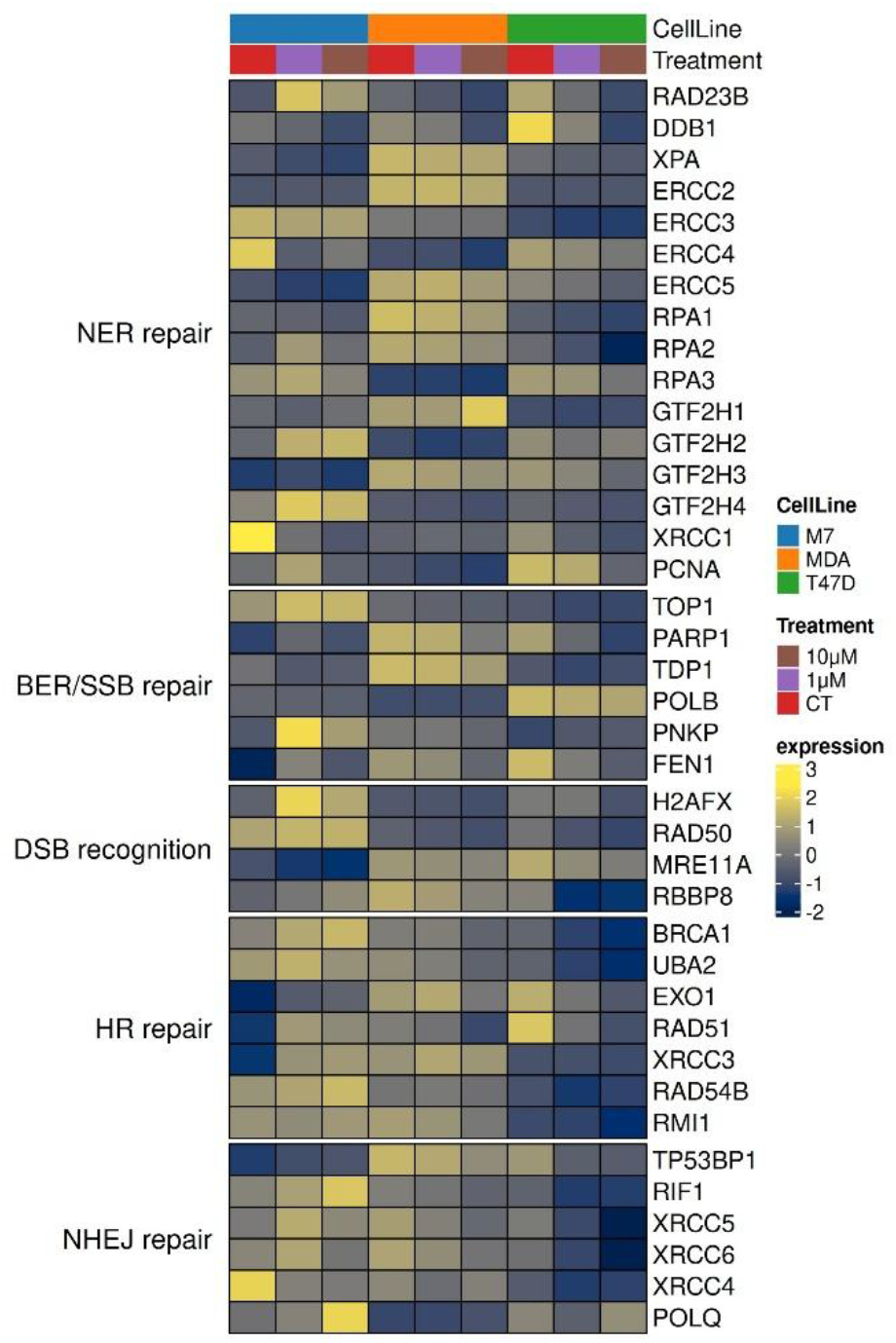
Heatmap of p53-independent genes involved in DNA repair. RNA expression levels of genes NOT on the direct p53-TGs involved in DNA repair. The three cell lines MCF-7 (WT p53), MDA-MB231 (p53 R280K) and T47D (L194F) were untreated or treated with 4NQO at 1µM or 10µM for 2h before RNA extraction.

**Fig. S4.**
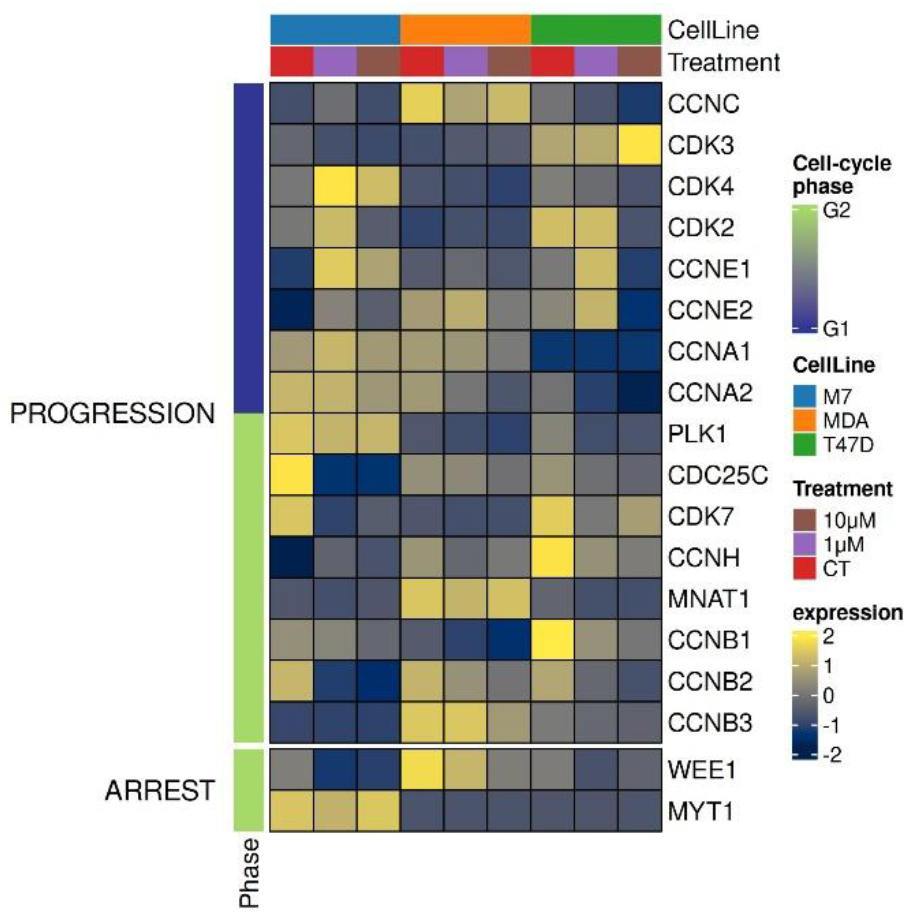
Heatmap of p53-independent genes involved in cell cycle control. RNA expression levels of genes NOT on the direct p53-TGs involved in DNA repair. The three cell lines MCF-7 (WT p53), MDA-MB231 (p53 R280K) and T47D (L194F) were untreated or treated with 4NQO at 1µM or 10µM for 2h before RNA extraction.

**Fig. S5.**
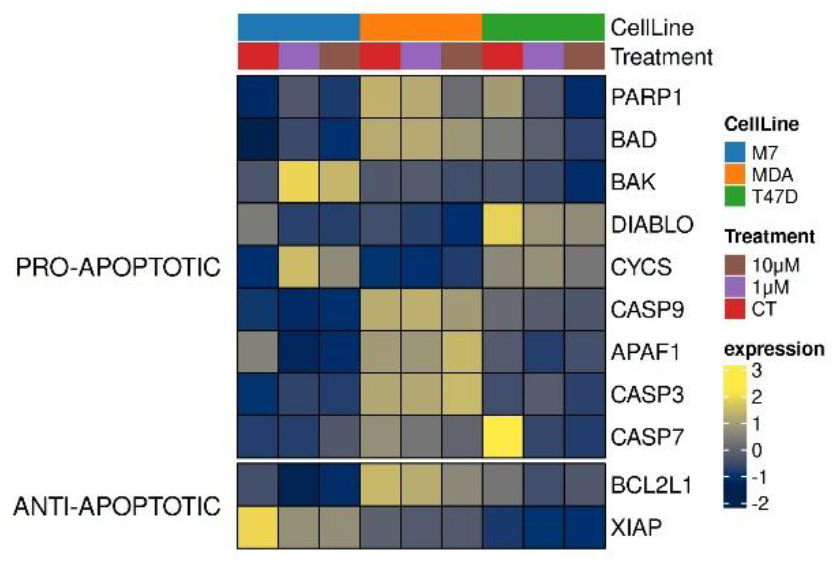
Heatmap of p53-independent genes involved in apoptotic cell death. RNA expression levels of genes NOT on the direct p53-TGs involved in DNA repair. The three cell lines MCF-7 (WT p53), MDA-MB231 (p53 R280K) and T47D (L194F) were untreated or treated with 4NQO at 1µM or 10µM for 2h before RNA extraction.

